# Conditional Protein Structure Generation with Protpardelle-1C

**DOI:** 10.1101/2025.08.18.670959

**Authors:** Tianyu Lu, Richard Shuai, Petr Kouba, Zhaoyang Li, Yilin Chen, Akio Shirali, Jinho Kim, Po-Ssu Huang

## Abstract

We present Protpardelle-1c, a collection of protein structure generative models with robust motif scaffolding and support for multi-chain complex generation under hotspot-conditioning. Enabling sidechain-conditioning to a backbone-only model increased Protpardelle-1c’s MotifBench score from 4.97 to 28.16, outperforming RFdiffusion’s 21.27. The crop-conditional all-atom model achieved 208 unique solutions on the La-Proteina all-atom motif scaffolding benchmark, on par with La-Proteina while having ∼10 times fewer parameters. At 22M parameters, Protpardelle-1c enables rapid sampling, taking 40 minutes to sample all 3000 MotifBench backbones on an NVIDIA A100-80GB, compared to 31 hours for RFdiffusion.

## 1 Introduction

To engineer proteins with function, it is often convenient to treat certain regions of a protein independently from the rest. Motifs are often defined as sets of amino acid patterns or conformations associated with specific functional sites. The ability to generate diverse scaffolds conditioned on a desired motif geometry is a key challenge in protein design. Motif scaffolding offers an approach to design functional *de novo* proteins if the motif geometry specification is accurately presented. The objective is to generate scaffolds that can host the motif in the desired and functionally relevant geometry, given a complete or partial functional motif. Conditional sampling allows the motif to be present in all the generated samples, in contrast to the traditional method of searching for motifs in a scaffold library [1, 2, 3].

Evaluation of a motif scaffolding model requires experimental data, but in the model development phase, we can benchmark the performance *in silico* with protein structure prediction models such as ESMFold [4] and count the number of unique samples that recapitulate the motif structure in the predicted structure. We consider two benchmarks of motif scaffolding, MotifBench [5] and RFdiffusion/La-Proteina [6, 7], spanning a diverse set of motif geometries from one to eight segments. Here, we describe modifications to the training dataset and the model architecture of Protpardelle [8] which enables the updated Protpardelle-1c `cc58` model to solve 22 of the 30 MotifBench problems with 164 total unique solutions. As a reference, RFdiffusion can solve 16 of the problems with 192 total unique solutions. The same changes also improve the Protpardelle-1c `cc91` all-atom model performance on the RFdiffusion/La-Proteina motif scaffolding benchmark, solving 22*/*26 tasks with 208 unique solutions. The list of available Protpardelle-1c models is described in Table 2. Code and examples are available at https://github.com/ProteinDesignLab/protpardelle-1c/tree/main.

## 2 Methods

Protpardelle is a diffusion model following the EDM framework that introduces the first all-atom generative design capability [8, 9]. In contrast to most other generative models of protein structure, Protpardelle is not equivariant to rotations and translations by design. As such, during training, the input is randomly rotated by a uniform sample from *SO(3)* and randomly translated by a sample from 𝒩 (0, 1). During sampling, the initial Gaussian noise is *SO(3)* invariant and the model picks a frame during the denoising process.

Conditional sampling with a diffusion model can be achieved with a combination of approaches, which we refer to the framework introduced in Didi *et al*. for details [10]. Here, we highlight reconstruction/classifier-based guidance and classifier-free guidance. The initial-release Protpardelle model was trained unconditionally and is only amenable to reconstruction guidance, in which at every denoising step, the negative gradient of a motif reconstruction loss is added to the score:

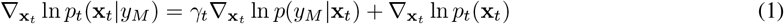

where **x**_*t*_ is the structure at denoising step *t* ∈ [0, 1], *p*_*t*_ is the model distribution, and *y*_*M*_ is the motif coordinates, and *p*(*y*_*M*_ | **x**_*t*_), i.e. the classifier term, is estimated with the mean squared error of the motif coordinates in **x**_*t*_ to the desired motif coordinates *y*_*M*_, where we assume the motif indices are known, i.e. given by the user or sampled from scaffold length range constraints.

However, this approach is often brittle as the guidance scale *γ*_*t*_ must be tuned empirically. In practice, setting *γ*_*t*_ too large risks broken chains where the motif is disconnected, while setting *γ*_*t*_ too small risks poor motif reconstruction. The search space for this hyperparameter is large, as it can also be time-dependent instead of a constant.

In classifier-free guidance, we amortize training of an unconditional and a conditional model into one model, by randomly dropping out the conditioning input some percentage of steps, here 5%. We can draw conditional samples with

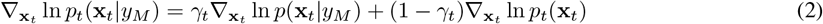

where the reversal in the *p*(**x**_*t*_ | *y*_*M*_) term is possible as the model is trained to attend to motif coordinates *y*_*M*_ as conditional input. The ln *p*_*t*_(**x**_*t*_) term is computed by setting the motif input to zeros. In all our experiments, we set *γ*_*t*_ = 1.

### 2.1 Augmented Datasets

To train Protpardelle-1c models, we curated augmented versions of the CATH dataset used to train previous versions of Protpardelle [11, 12].

#### 2.1.1 AI-CATH: Sequence design augmented dataset

The augmented Ingraham CATH (AI-CATH) dataset contains the designable ESMFold predicted structures of 32 ProteinMPNN redesigned sequences for each CATH structure belonging to the original CATH training dataset [4, 13, 11]. Similar to MultiFlow [14], we only train on self-consistent structures (scRMSD < 2.0 Å and pLDDT *>* 80), arriving at a dataset of 337,936 structures.

#### 2.1.2 MD-CATH: Molecular dynamics augmented CATH

We curated a dataset of protein conformational ensembles sampled from the dataset of molecular dynamics (MD) simulations MD-CATH [15]. The MD-CATH dataset provides MD simulations of 5398 protein domains from CATH 4.2.0 dataset which passed the criteria for simulations such as length between 50 and 500 residues or absence of gaps and non-standard AAs. The average trajectory length is 464 ns, with frames saved every nanosecond. We used only the version of mdCATH simulated at 320K as it is closer to physiological temperatures than the other versions. At this temperature there are 5 trajectories for each protein. For each protein, we subsample 32 conformations from the MD ensemble and we perform Rosetta cartesian minimization for the sampled conformations to correct MD artifacts. Details on the sub-sampling and minimization are given in Appendix B.

## 3 Results

### 3.1 Unconditional Coverage

We verified the coverage of our baseline unconditional model `bb81` and unconditional samples from the motif scaffolding model `cc58` with SHAPES [16]. Both models adequately cover the CATH structure distribution (Figure 1) and we proceeded with evaluations of conditional generation described below.

**Figure 1:**
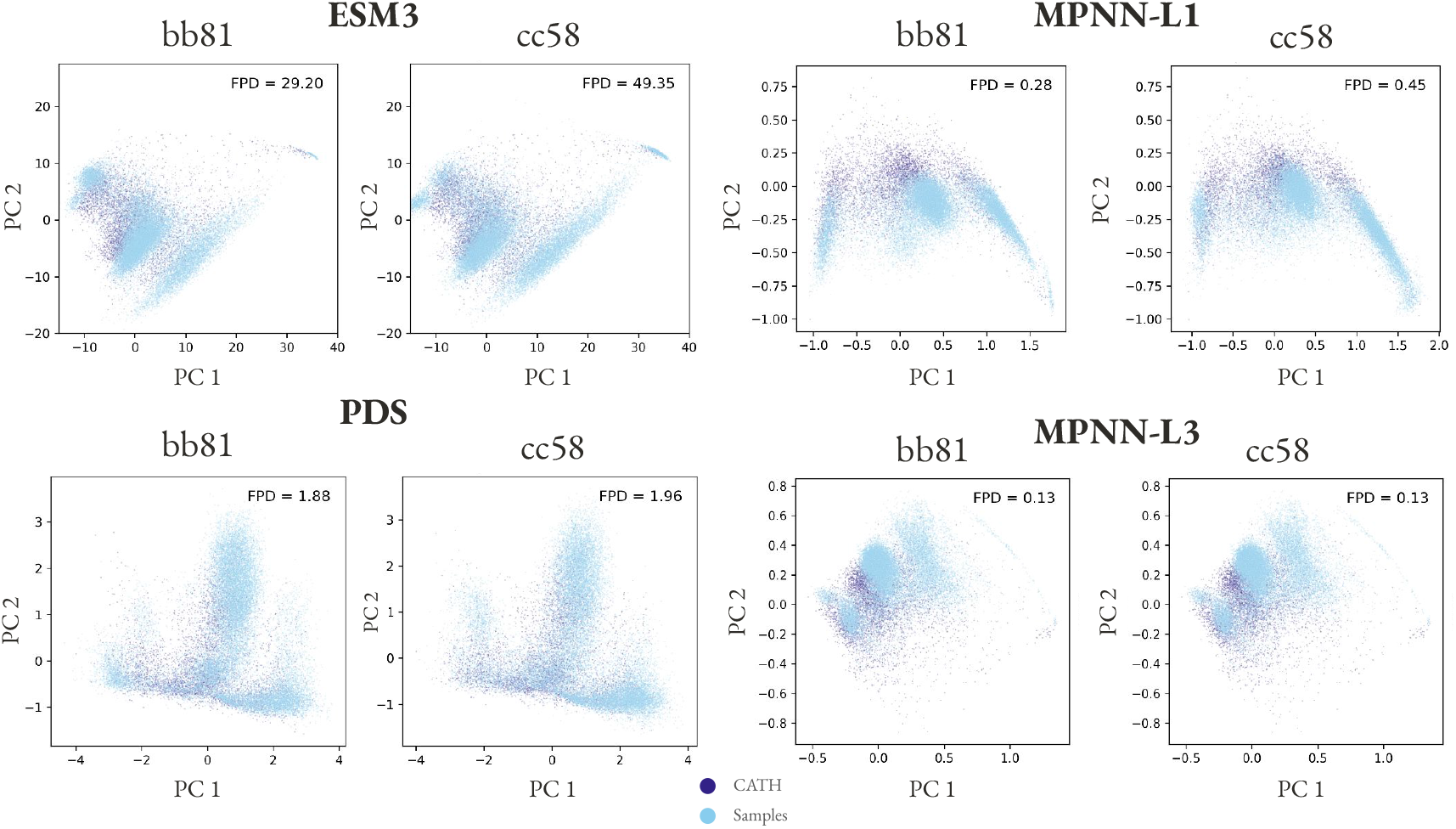
Protpardelle-1c samples cover the CATH distribution. ∼ 20k samples were drawn from each model (`bb81` and `cc58`) and embeddings for the samples and CATH structures were computed with ESM3, Protein Domain Segmentor (PDS), and ProteinMPNN encoder layers 1 and 3 (MPNN-L1 and MPNN-L3). We follow the CATH filter in SHAPES: keep structures with resolution < 3.0 Å, Rfree < 0.25, and no unresolved residues [16]. The first two principal components of the embeddings are shown. The Fréchet Protein Distance (FPD) denotes the coverage where lower is better.

### 3.2 MotifBench

With the initial-release Protpardelle model, we achieved a MotifBench score of 4.97 using the best hyperparameters of `step_scale` = 1.1 and *γ*_*t*_ = 0.1 with replacement guidance. Given that we used a backbone-only model, the guidance term in Equation 1 was independent of motif sidechain atom coordinates. This is not optimal as polar residues may become buried and hydrophobic residues may become exposed in the sampled scaffolds, leading to lower success rates. To test this hypothesis, during ProteinMPNN sequence redesign, we allowed all positions, including the motif residues, to be redesigned, finding that indeed the model achieved a higher MotifBench score of 10.18. We reasoned that a model that can be conditioned on both backbone and sidechain coordinates of the motif would simultaneously address the brittleness of reconstruction guidance and find scaffolds which can better host the motif sidechains.

We trained a crop-conditional motif scaffolding model, `cc58`, identical to what was described in the original Protpardelle [8] and analogous to the Doob’s *h*-transform conditional training setup in Algorithm 5 of Didi *et al*. [10]. We concatenate the motif coordinates channel-wise such that a model input with dimensions *B, L, N*, 2 × *X* becomes *B, L, N*, 3 × *X*, where *B* is the batch size, *L* is the sequence length, *N* = 37 is the number of unique PDB atom types per residue, and *X* = 3 is the *x, y, z* coordinates. The 2× allows for self-conditioning (6 channels), thus expanding it to 3× allows for both self-conditioning and motif-conditioning (9 channels). A limitation is that the motif indices must be pre-specified. We leave concatenation along the *L* dimension, which would allow unindexed motifs as in RFdiffusion2 and La-Proteina, for future work [17, 7]. We change the location of where the noise residual is added to match the architecture of the adaptive layer normalization setup in DiT [18]. This change reduced the trainable parameters from 33M to 22M since the previous architecture has the noise residual added at the feedforward layer where the dimensionality expands 4 ×, causing an excessive number of parameters (11M) to encode the noise level. Other details of the model architecture and the training noise schedule remain unchanged.

The backbone-only motif scaffolding model, `cc58`, was trained for 4.38M steps with an NVIDIA A100-80GB on AI-CATH. The initial-release Protpardelle model was trained with rotary positional encoding that used raw tensor indices, such that gaps in protein structures due to unresolved residues were not correctly considered, i.e. the model treats the two flanking residues of a gap to be adjacent. While it appears that we can resolve this issue by deriving the positional encoding from PDB residue indices, the PDB files used in the CATH dataset contain inconsistencies in residue indexing that cause gap lengths to be inconsistent with the actual number of unresolved residues when compared to the `SEQRES` sequence, e.g. PDB: 1914. We resolved such residue indexing inconsistencies in the AI-CATH dataset. We note that as the PDB file format is deprecated, deriving positional encodings from the `.cif` file’s `_atom_site.label_seq_id` indices would correct indexing inconsistencies. During training, motifs are randomly generated as crops of the input structure with the following parameters: contiguous (5%), discontiguous (90%), sidechain conditioned (90%), maximum contiguous segment length (12), maximum number of discontiguous residues forming the motif (24), maximum distance between residue pairs forming the motif (45.0 Å), and translating the structure such that the motif coordinates have center of mass at the origin, before the default rotation/translation augmentations and masking non-existent motif atom coordinates to zeros.

A solution to a MotifBench problem must satisfy the following criteria using ESMFold-predicted structures of 8 ProteinMPNN sequences per scaffold, fixing the (possibly a subset of) motif amino acid types:

1. Motif N, C*α*, and C RMSD < 1.0 Å.
2. Scaffold C*α*-only RMSD < 2.0 Å.

The number of unique solutions is the number of clusters with TM score threshold of 0.5 obtained with `Foldseek-Cluster`. Protpardelle-1c is able to solve the additional problems `13_4JHW`, `21_1B73`, `23_1MPY`, `24_1QY3`, `25_2RKX`, `26_3B5V`, `27_4XOJ`, and `29_6CPA`, offering solutions to previously difficult multi-segment tasks (Figure 2). No solutions were found for `01_1LDB`, `03_2CGA`, `08_7AD5`, `12_4JHW`, `18_7MQQ`, `19_7MQQ`, `20_7UWL`, and `30_7UWL`. Across runs with different random seeds, we occasionally observe solutions for these motifs though the MotifBench score remains fairly consistent. We also confirm that the PDBs for all MotifBench problems for which Protpardelle-1c finds at least one solution are not present in the training data. We highlight four examples of successful scaffolds in Figure 3. Note that the highlighted solution for `1QY3` has 9 strands instead of the typical 11 strand barrel found in natural GFP. Although having fewer total unique solutions than RFdiffusion, the MotifBench score is higher for Protpardelle-1c due to additional solutions to several previously unsolved problems, while the extra RFdiffusion solutions are concentrated mainly on `07_6E6R`.

**Figure 2:**
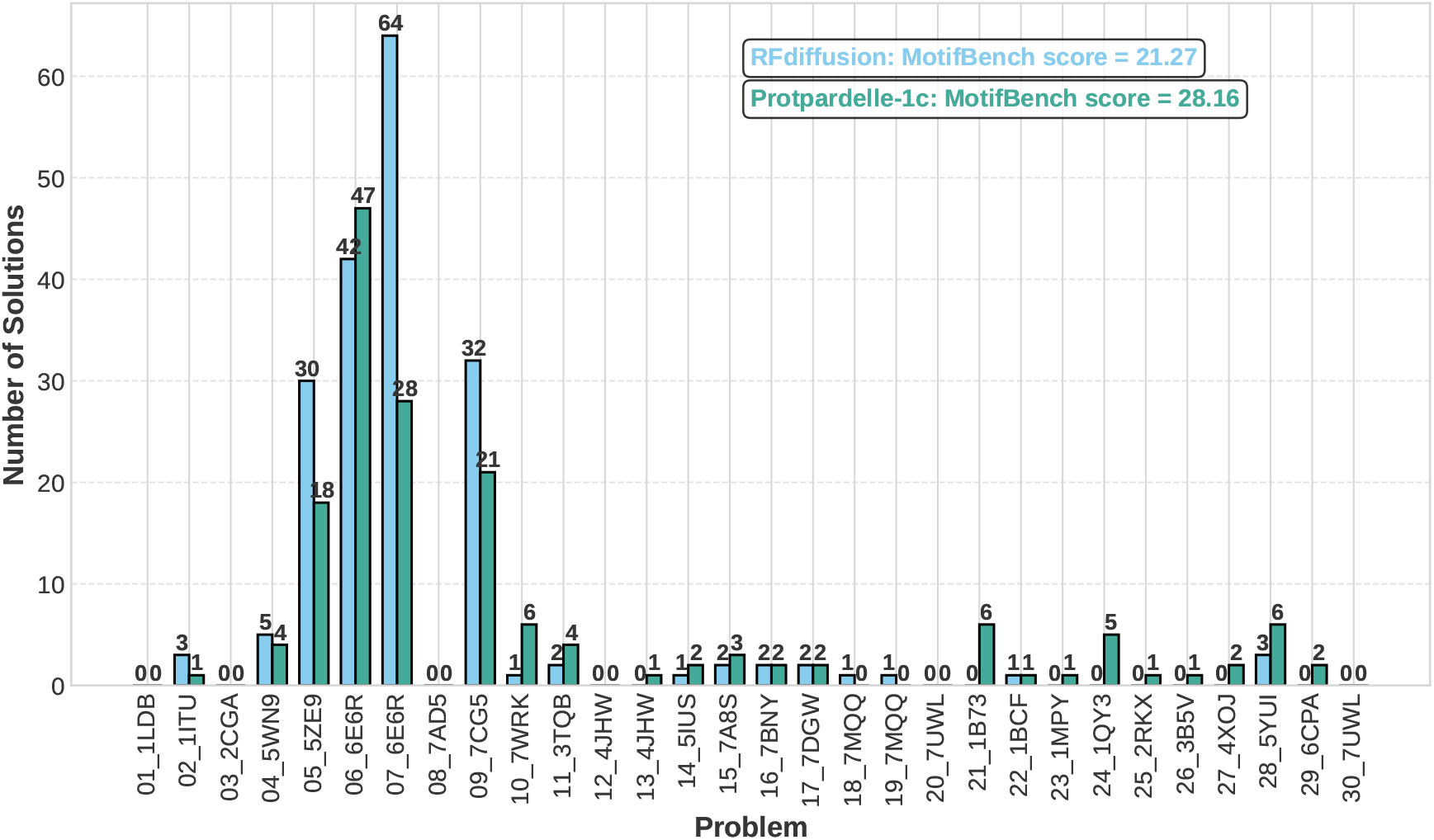
Protpardelle-1c outperforms RFdiffusion on MotifBench. Per-problem unique successes on MotifBench of Protpardelle-1c `cc58` compared to RFdiffusion.

**Figure 3:**
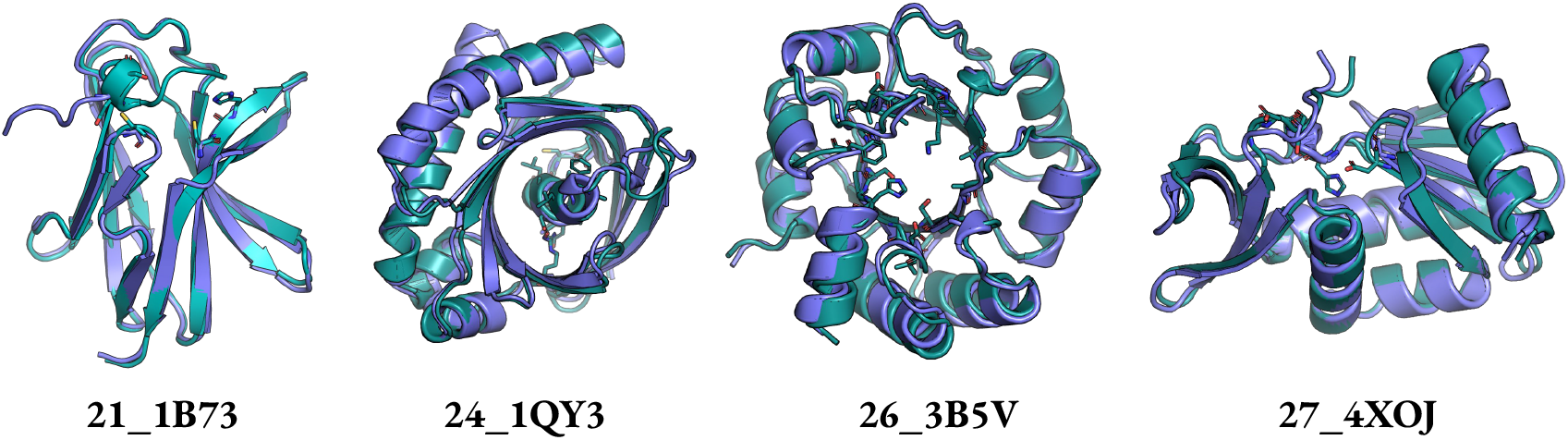
Examples of MotifBench solutions on previously unsolved cases. `21_1B73`: Glutamate racemase. `24_1QY3`: GFP fluorophore. `26_3B5V`: Retroaldolase. `27_4XOJ`: Trypsin. Purple: sampled structure. Teal: ESMFold-predicted structure. Motif atoms of the ESMFold-predicted structures are shown in licorice. All structures shown pass MotifBench criteria.

### 3.3 RFdiffusion and La-Proteina Motif Scaffolding Benchmark

We benchmark the backbone-only model `cc58` and the all-atom model `cc91` against motifs scaffolding tasks first introduced in RFdiffusion [6] and modified by La-Proteina [7]. A solution must satisfy the following criteria:

1. Motif C*α* RMSD < 1 Å.
2. Motif all-atom RMSD < 2 Å.
3. All-atom scRMSD < 2 Å.

The number of unique solutions is the number of clusters with TM score threshold of 0.5 obtained with `Foldseek-Cluster`. Since the backbone-only model `cc58` does not directly generate sidechains, we cannot compute criteria 2 and 3 and compare with La-Proteina directly. Instead, we replace criterion 2 with the motif all-atom scRMSD between the coordinates of the ground truth motif and the ESMFold-predicted structure and replace criterion 3 with the C*α* scRMSD between the sample and the ESMFold-predicted structure. The all-atom model `cc91`‘s evaluation criteria remain unchanged. For motifs with unresolved atoms, e.g. `5TRV`, we do not remodel the missing motif atoms and compute the all-atom RMSD only on the resolved atoms.

Previously, the original all-atom Protpardelle model solved 4*/*26 tasks with a total of 4 unique solutions whereas the Protpardelle-1c all-atom `cc91` model solved 22*/*26 tasks with a total of 208 unique solutions (Figure 4, Table 1). The backbone-only model `cc58` solved 26*/*26 tasks with a total of 787 unique solutions with MPNN-8 (Table 3). MPNN-X refers to sampling X ProteinMPNN sequences allows per scaffold and a scaffold is considered a success if at least one design passes all criteria. A more comparable setting is with MPNN-1 as La-Proteina evaluates a single sequence per scaffold. In this setting, `cc58` solved 25*/*26 tasks with 424 unique solutions. Motif all-atom scRMSD < 2 Å may be too lenient both with respect to the often sub-Ångstrom precision required for motifs to be functional and that fewer residues are involved in the all-atom scRMSD calculation compared to La-Proteina where the whole protein is considered. Thus, we also report the success rate with the strict criterion of all-atom motif scRMSD < 1 Å in Table 4 (together with motif C*α* RMSD < 1 Å and scaffold C*α* scRMSD < 2 Å). With the strict criterion, `cc58` solved 14*/*26 tasks with 295 unique solutions with MPNN-8 and solved 10/26 problems with 107 unique solutions with MPNN-1. While a direct comparison between the backbone-only `cc58` model and the all-atom models (La-Proteina and `cc91`) is not possible due to the different success criteria, we observe that `cc91`‘s performance is roughly on par with the performance of `cc58` MPNN-1, which matches the single ProteinMPNN sequence used in the final step of `cc91`‘s denoising trajectory. Examples of successes are shown in Figure 5.

**Table 1:** La-Proteina motif scaffolding benchmark: La-Proteina and cc91.

| Motif | # segments | La-Proteina (indexed) |  | La-Proteina (unindexed) |  | cc91 |  |
| --- | --- | --- | --- | --- | --- | --- | --- |
|  |  | All | Unique | All | Unique | All | Unique |
| 1ycr | 1 | 123 | 38 | 120 | 68 | 65 | 34 |
| 7mrx_128 | 1 | 22 | 17 | 86 | 36 | 9 | 6 |
| 5trv_long | 1 | 91 | 9 | 26 | 14 | 21 | 21 |
| 5tpn | 1 | 55 | 1 | 34 | 13 | 15 | 10 |
| 4zyp | 1 | 11 | 2 | 82 | 7 | 2 | 1 |
| 7mrx_85 | 1 | 16 | 4 | 104 | 6 | 18 | 7 |
| 5trv_med | 1 | 65 | 3 | 15 | 6 | 9 | 8 |
| 3ixt | 1 | 34 | 6 | 50 | 5 | 3 | 3 |
| 7mrx_60 | 1 | 7 | 3 | 73 | 3 | 8 | 3 |
| 5trv_short | 1 | 5 | 1 | 2 | 2 | 0 | 0 |
| 6e6r_short | 1 | 35 | 8 | 0 | 0 | 3 | 2 |
| 6e6r_med | 1 | 73 | 22 | 0 | 0 | 15 | 14 |
| 6e6r_long | 1 | 71 | 43 | 0 | 0 | 16 | 16 |
| 5wn9 | 1 | 0 | 0 | 0 | 0 | 0 | 0 |
| 1prw | 2 | 175 | 20 | 122 | 11 | 161 | 20 |
| 6vw1 | 2 | 21 | 1 | 60 | 6 | 14 | 5 |
| 2kl8 | 2 | 165 | 1 | 156 | 1 | 176 | 1 |
| 5ius | 2 | 16 | 2 | 1 | 1 | 0 | 0 |
| 4jhw | 2 | 2 | 1 | 0 | 0 | 0 | 0 |
| 1qjg_native | 3 | 72 | 13 | 76 | 54 | 23 | 17 |
| 7k4v | 3 | 116 | 11 | 35 | 35 | 14 | 2 |
| 1qjg | 3 | 72 | 4 | 58 | 23 | 16 | 13 |
| 5yui | 3 | 11 | 4 | 21 | 13 | 3 | 3 |
| 5aou | 3 | 145 | 4 | 9 | 9 | 91 | 7 |
| 5aou_quad | 4 | 171 | 2 | 92 | 37 | 56 | 6 |
| 1bcf | 4 | 189 | 7 | 148 | 7 | 196 | 9 |
| **Total** |  | **1763** | **227** | **1370** | **357** | **934** | **208** |

**Table 2:** Available Protpardelle-1c models. Notes: **bb81_epoch450**: Unconditional model trained on AI-CATH; **bbmd_epoch500**: Unconditional model trained on MD-CATH; **cc58_epoch416**: MotifBench benchmark model; **cc58_epochX**: Additional checkpoints of cc58: 521, 595, 649, 777, and 838; **cc58-minimpnn**: Trained on cc58_epoch595 1-step x0 predicted structures; **cc78_epoch1431**: Experimental: residue indices are tied across chains, favors homodimers; **cc83_epoch2616**: BindCraft benchmark model; **cc89_epoch415**: Sequence must be provided at all sampling steps; **cc91_epoch383**: Allatom model trained on AI-CATH; **cc91_tip_epoch480**: cc91 finetuned on sidechain tip atom conditioning task; **cc94_epoch3100**: cc91 finetuned on multi-chain data but no hotspot; **cc95_epoch3490**: cc83 finetuned with heavier hotspot dropout.

| Name | Monomers | Multi-chain | Model Type | Positional Encoding |
| --- | --- | --- | --- | --- |
| **bb81_epoch450** | 1 | 0 | Backbone | Rotary |
| **bbmd_epoch500** | 1 | 0 | Backbone | Rotary |
| **cc58_epoch416** | 1 | 0 | Backbone | Rotary |
| **cc58_epochX** | 1 | 0 | Backbone | Rotary |
| **cc58-minimpnn** | 1 | 0 | Sequence design | - |
| **cc78_epoch1431** | 0 | 1 | Backbone | Relative + Relchain |
| **cc83_epoch2616** | 0.5 | 0.5 | Backbone | Relative |
| **cc89_epoch415** | 1 | 0 | Allatom Sequence Mask | Rotary |
| **cc91_epoch383** | 1 | 0 | Allatom No Mask | Relative |
| **cc91_tip_epoch480** | 1 | 0 | Allatom No Mask | Relative |
| **cc94_epoch3100** | 0.5 | 0.5 | Allatom No Mask | Relative |
| **cc95_epoch3490** | 0.5 | 0.5 | Backbone | Relative + Relchain |

**Table 3:** La-Proteina motif scaffolding benchmark: cc58.

| Motif | # segments | cc58 MPNN-1 |  | cc58 MPNN-4 |  | cc58 MPNN-8 |  |
| --- | --- | --- | --- | --- | --- | --- | --- |
|  |  | All | Unique | All | Unique | All | Unique |
| 1ycr | 1 | 128 | 69 | 173 | 88 | 189 | 105 |
| 7mrx_128 | 1 | 14 | 10 | 39 | 21 | 57 | 26 |
| 5trv_long | 1 | 25 | 14 | 53 | 25 | 73 | 30 |
| 5tpn | 1 | 87 | 30 | 132 | 46 | 150 | 47 |
| 4zyp | 1 | 19 | 6 | 51 | 15 | 80 | 20 |
| 7mrx_85 | 1 | 46 | 13 | 98 | 25 | 119 | 27 |
| 5trv_med | 1 | 18 | 10 | 38 | 16 | 56 | 22 |
| 3ixt | 1 | 25 | 9 | 54 | 17 | 69 | 18 |
| 7mrx_60 | 1 | 42 | 13 | 81 | 13 | 120 | 26 |
| 5trv_short | 1 | 3 | 1 | 13 | 3 | 11 | 4 |
| 6e6r_short | 1 | 35 | 24 | 67 | 52 | 95 | 63 |
| 6e6r_med | 1 | 88 | 62 | 144 | 102 | 166 | 112 |
| 6e6r_long | 1 | 84 | 55 | 143 | 80 | 165 | 101 |
| 5wn9 | 1 | 0 | 0 | 2 | 2 | 1 | 1 |
| 1prw | 2 | 176 | 22 | 191 | 26 | 198 | 18 |
| 6vw1 | 2 | 47 | 5 | 87 | 9 | 102 | 5 |
| 2kl8 | 2 | 190 | 1 | 200 | 1 | 199 | 1 |
| 5ius | 2 | 3 | 1 | 8 | 3 | 14 | 4 |
| 4jhw | 2 | 2 | 2 | 2 | 1 | 7 | 7 |
| 1qjg_native | 3 | 52 | 23 | 92 | 34 | 124 | 43 |
| 7k4v | 3 | 26 | 4 | 53 | 3 | 88 | 6 |
| 1qjg | 3 | 48 | 26 | 85 | 40 | 124 | 62 |
| 5yui | 3 | 2 | 2 | 9 | 8 | 14 | 11 |
| 5aou | 3 | 66 | 9 | 128 | 6 | 147 | 10 |
| 5aou_quad | 4 | 57 | 5 | 119 | 6 | 146 | 8 |
| 1bcf | 4 | 186 | 8 | 200 | 4 | 200 | 10 |
| **Total** |  | **1469** | **424** | **2262** | **646** | **2714** | **787** |

**Table 4:** La-Proteina motif scaffolding benchmark: cc58 with all-atom motif scRMSD < 1 Å instead of 2 Å.

| Motif | # segments | cc58 MPNN-1 Strict |  | cc58 MPNN-4 Strict |  | cc58 MPNN-8 Strict |  |
| --- | --- | --- | --- | --- | --- | --- | --- |
|  |  | All | Unique | All | Unique | All | Unique |
| 1ycr | 1 | 48 | 30 | 85 | 45 | 120 | 64 |
| 7mrx_128 | 1 | 0 | 0 | 0 | 0 | 0 | 0 |
| 5trv_long | 1 | 0 | 0 | 0 | 0 | 0 | 0 |
| 5tpn | 1 | 0 | 0 | 1 | 1 | 1 | 1 |
| 4zyp | 1 | 0 | 0 | 0 | 0 | 0 | 0 |
| 7mrx_85 | 1 | 0 | 0 | 0 | 0 | 0 | 0 |
| 5trv_med | 1 | 0 | 0 | 0 | 0 | 0 | 0 |
| 3ixt | 1 | 0 | 0 | 20 | 6 | 19 | 6 |
| 7mrx_60 | 1 | 0 | 0 | 0 | 0 | 0 | 0 |
| 5trv_short | 1 | 0 | 0 | 0 | 0 | 0 | 0 |
| 6e6r_short | 1 | 3 | 3 | 25 | 15 | 23 | 16 |
| 6e6r_med | 1 | 20 | 18 | 61 | 43 | 79 | 59 |
| 6e6r_long | 1 | 28 | 23 | 66 | 43 | 92 | 60 |
| 5wn9 | 1 | 0 | 0 | 0 | 0 | 2 | 2 |
| 1prw | 2 | 163 | 12 | 187 | 24 | 194 | 24 |
| 6vw1 | 2 | 0 | 0 | 0 | 0 | 0 | 0 |
| 2kl8 | 2 | 0 | 0 | 0 | 0 | 0 | 0 |
| 5ius | 2 | 0 | 0 | 0 | 0 | 0 | 0 |
| 4jhw | 2 | 0 | 0 | 0 | 0 | 0 | 0 |
| 1qjg_native | 3 | 12 | 9 | 33 | 13 | 57 | 25 |
| 7k4v | 3 | 1 | 1 | 5 | 2 | 1 | 1 |
| 1qjg | 3 | 11 | 8 | 36 | 16 | 52 | 22 |
| 5yui | 3 | 0 | 0 | 0 | 0 | 0 | 0 |
| 5aou | 3 | 6 | 1 | 19 | 2 | 21 | 5 |
| 5aou_quad | 4 | 0 | 0 | 0 | 0 | 2 | 1 |
| 1bcf | 4 | 71 | 2 | 137 | 9 | 168 | 9 |
| **Total** |  | **363** | **107** | **675** | **219** | **831** | **295** |

**Figure 4:**
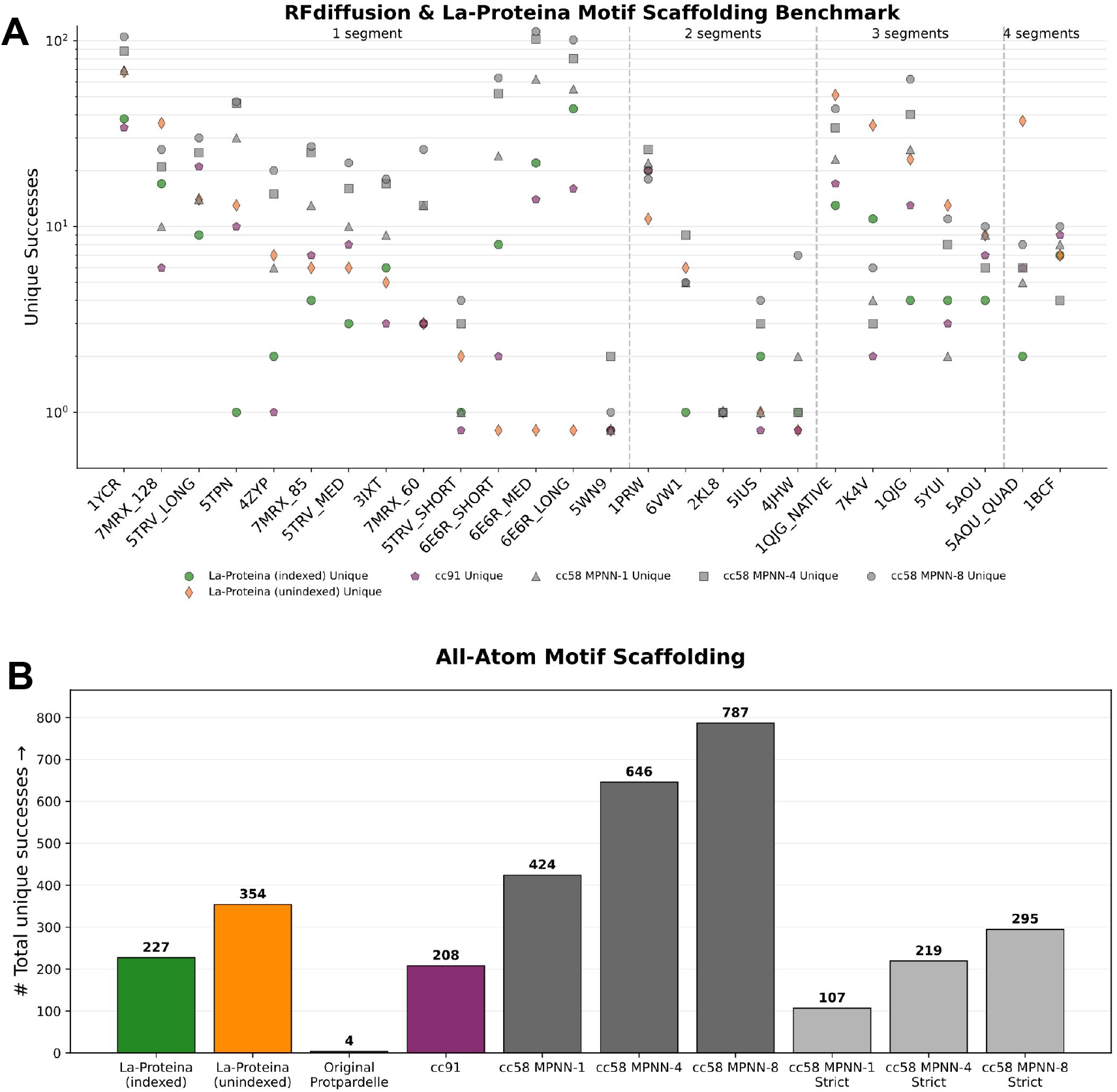
Protpardelle-1c is competitive with La-Proteina on all-atom motif scaffolding. Per-problem unique successes on motif scaffolding tasks initially introduced in RFdiffusion compared against La-Proteina. (A) Number of unique successes out of 200 samples per motif. Note that only `cc91` is directly comparable with La-Proteina due to the difference in the definition of success for the backbone-only `cc58` model. (B) Sum of unique successes out of 5200 total sampled scaffolds. Strict refers to motif all-atom scRMSD < 1 Å.

**Figure 5:**
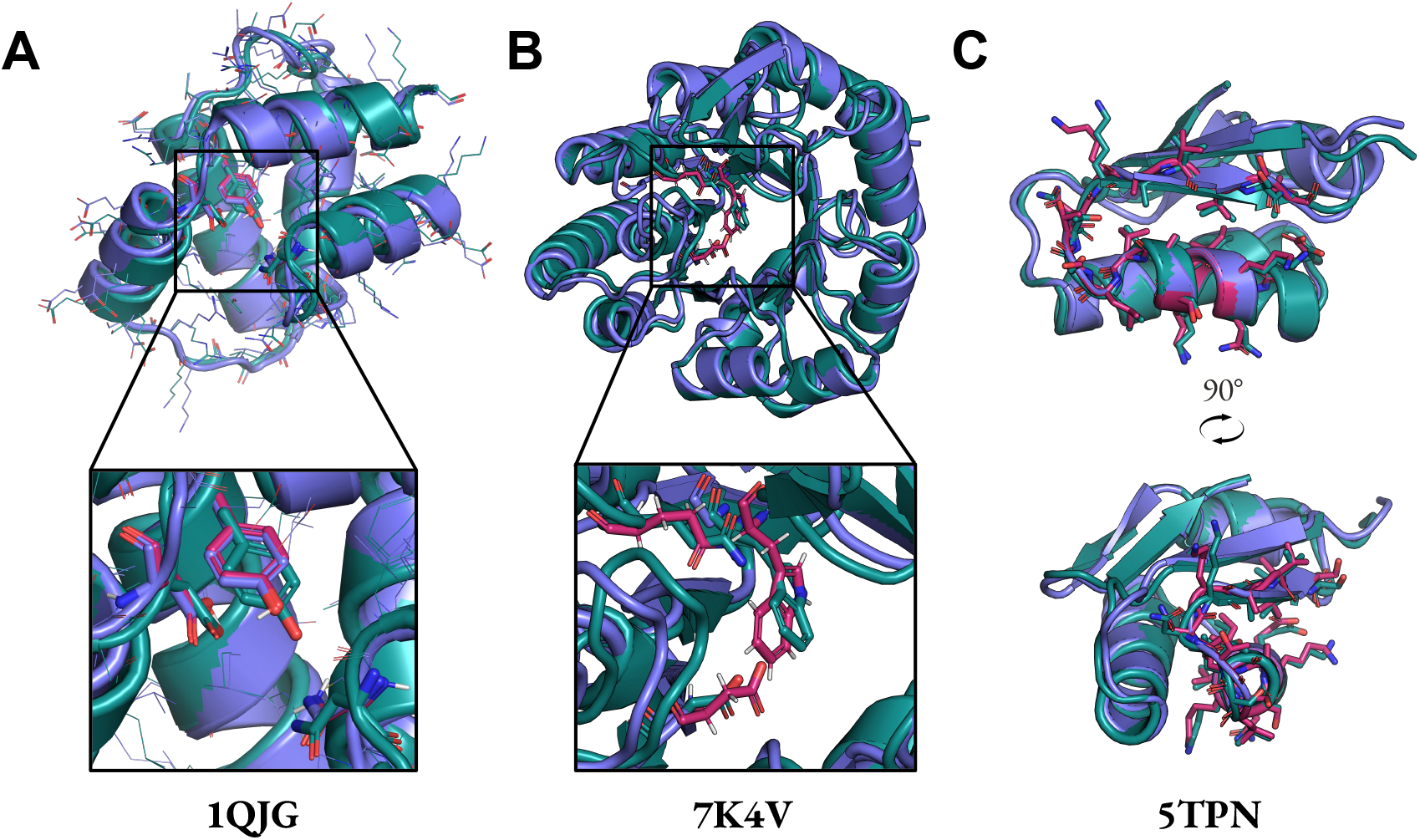
Examples of Protpardelle-1c scaffolds on RFdiffusion/La-Proteina motifs. In all panels, slate: sampled structure, deepteal: ESMFold predicted structure, warmpink: motif. (A) `cc91`. (B) `cc58` with all-atom motif scRMSD < 2 Å. (C) `cc58` with all-atom motif scRMSD < 1 Å.

### 3.4 Backbone-only vs. All-Atom

The backbone-only model denoises coordinates of the N, CA, C, O backbone atoms for each residue. The all-atom model denoises both backbone atoms and sidechain atoms. In the original Protpardelle, we define a superposition scheme in which for the first `skip_mpnn_proportion` denoising steps, the sequence is sampled from a uniform prior on all amino acid types, then up to but excluding the final step, the sequence is sampled from MiniMPNN, a noise-conditional ProteinMPNN model trained on the 1-step denoised predicted structure 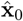. In Protpardelle-1c, we retrain MiniMPNN using `cc58-epoch416` as the pretrained structure denoiser. As MiniMPNN is not used until the backbone structure is roughly determined, we only train on the lower noise levels corresponding to the latter half of denoising. We only train on features derived from C*α* coordinates as the precise N, C*α*, C, O geometries are not yet formed when MiniMPNN is enabled. Following the original Protpardelle, we use full ProteinMPNN to design the final sequence. We call this process stage-1 sampling and use this to sample from `cc91`.

In Protpardelle-1c, we introduce a simpler all-atom sampling scheme which is distinguished from the superposition scheme by the name `uniform-steps`. We change the name of the original scheme to `jump-steps` to highlight the non-uniform denoising steps taken. In the `jump-steps` scheme, we cache the most recently observed **x**_*t*_ state for each amino acid type in an `atom73` format: backbone atoms plus all combinations of residue types and sidechain atom types. The next time an amino acid type is re-observed, the denoised output is computed in one step directly from the **x**_*t*_ state when the atom was previously observed. In the `uniform-steps` scheme, we take a step directly from the previous step **x**_*t* − 1_. In both schemes, fresh noise is injected in the dimensions corresponding to non-existent sidechain atoms, i.e. dummy atoms. This noise scales with the denoising schedule and is centered on the C*α* of each residue of the predicted denoised structure 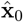 at the current step.

We modify the default stage-2 sampling protocol from the original Protpardelle to directly model the effect of sidechain-driven backbone conformation change. The default stage-2 sampling protocol in Protpardelle-1c is to apply all-atom partial diffusion to the output of stage-1 sampling. In contrast to the original Protpardelle where the backbone coordinates were fixed, we allow backbone flexibility to accommodate the final designed sequence from stage-1.

Finally, we also trained an all-atom model, `cc89`, in which the sequence is always provided to the model through the atom mask, where non-existent atoms are masked to be all zeros. The `cc89` model can be used for refinement and sidechain-aware partial diffusion. Given the same partial diffusion noise level, `cc89` samples a distinct space of structures, a CDR3 loop from 7EOW chain B in this example, compared to the loops sampled by an all-atom model where dummy atom coordinates are not masked (`cc94`) and backbone-only models trained on either AI-CATH or MD-CATH. At the same noise level of 150/500 rewind steps, `cc89` samples sidechain conformations with less fluctuation on the side of the CDR3 loop which packs against the nanobody framework (Figure 6A, right) where the solvent-exposed residues have more fluctuation (Figure 6A, left). In contrast, the other three models `cc94`, `bb81`, and `bbmd` do not have such residue-specific behavior: the fluctuations across the loop are more homogeneous than in `cc89` (Figure 6B, C, D) and more diffuse, as is evident in the projected CDR3 loop coordinates into the first two principal components (Figure 7).

**Figure 6:**
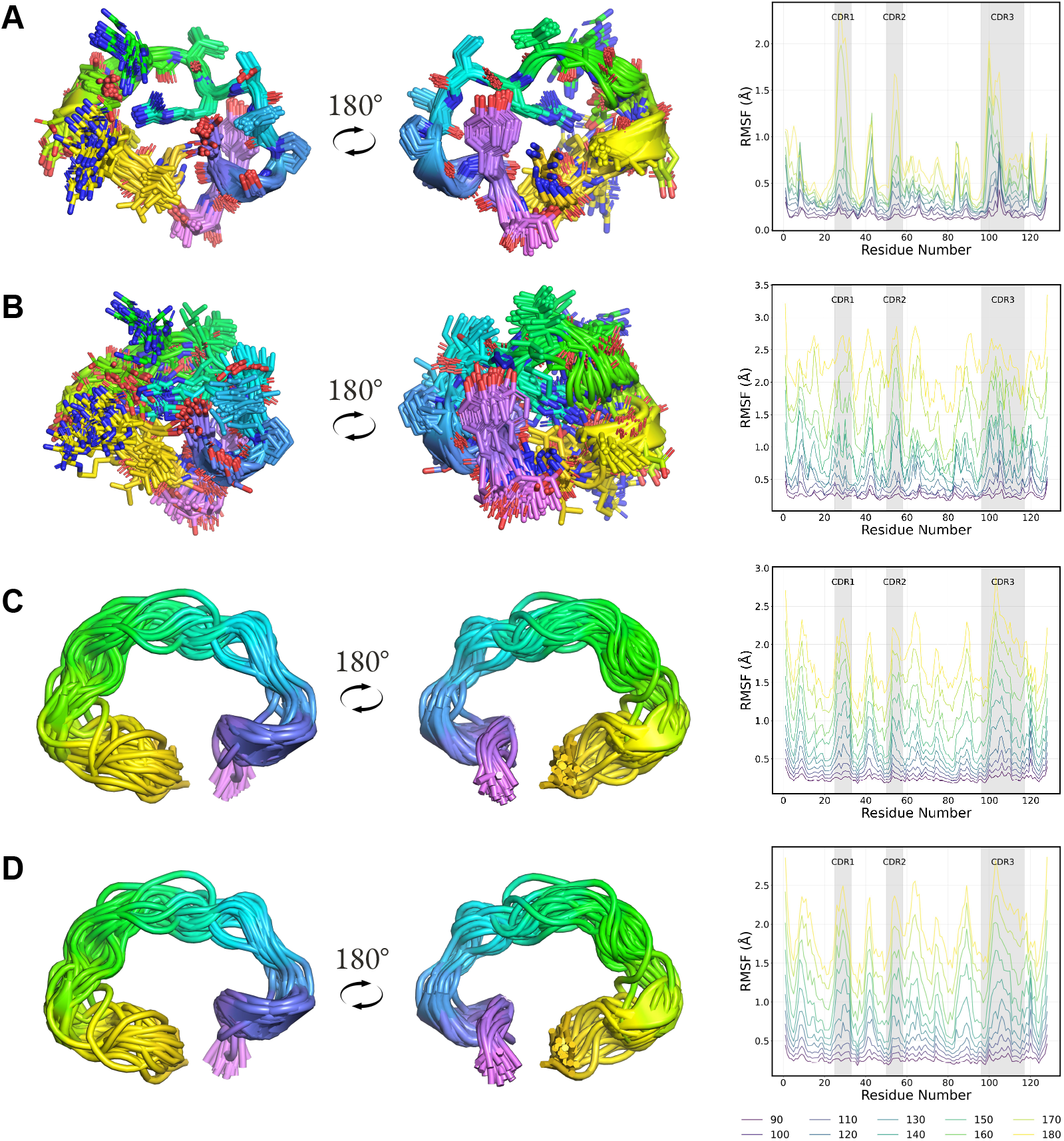
An all-atom model with sequence mask cc89 sample position-heterogeneous CDR3 loops. Partial diffusion rewind steps 150 out of 500 total denoising steps using (A) all-atom `cc89` model, (B) all-atom `cc94` model, (C) backbone-only `bb81` model, (D) backbone-only `bbmd` model. The per-residue Root Mean Square Fluctuation profiles are shown for each model at different partial diffusion rewind steps.

**Figure 7:**
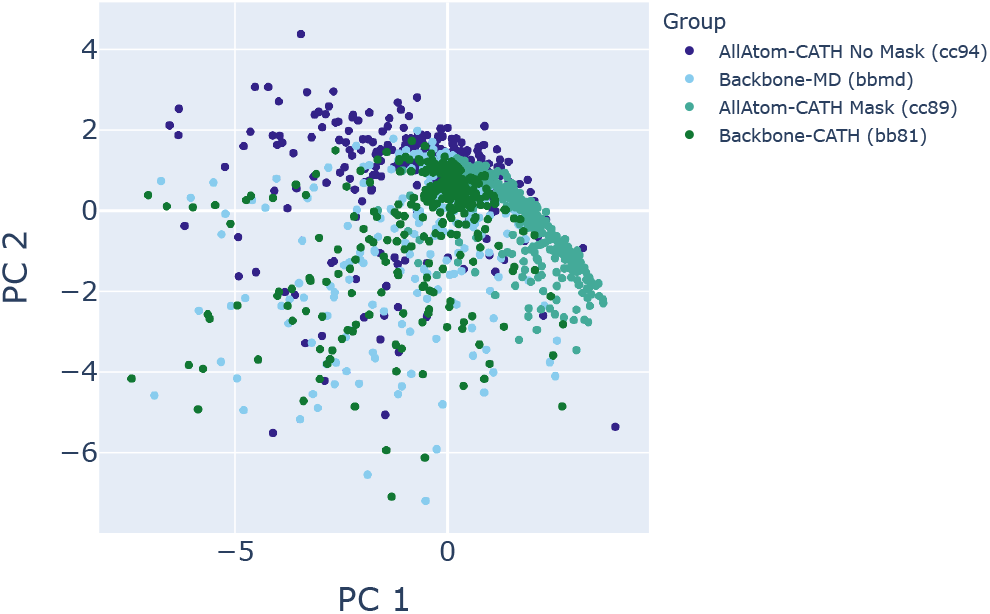
An all-atom model with sequence mask cc89 samples distict loop conformations. C*α* coordinates of the CDR3 loop are projected into their first two principal components. AllAtom-CATH No Mask: `cc94`, Backbone-MD: `bbmd`, AllAtom-CATH Mask: `cc89`, Backbone-CATH: `bb81`. The `cc89` model samples a more restricted space of loop conformations.

### 3.5 Multi-chain Conditional Generation

The binder design problem can be treated as a special case of the motif scaffolding problem, in which the target structure is the motif and the binder is the piece to generate. To handle multiple chains, we use learned relative positional embeddings added to the attention matrix analogous to AlphaFold2 [19], with a residue index gap between chains of 200. We train on a 50*/*50 mix of the single-chain augmented CATH dataset described previously and a curated chain pair dataset following Boltz-1 [20] (Appendix A). For multi-chain examples, we randomly select one chain as the target (motif) and take Unif(3, 8) closest residues to the binder chain as the hotspots, dropping it out 10% of the time. Paratope residues, defined as the residues on the binder chain used to choose the hotspots, are included as motif 50% of the time, allowing partial-paratope completions in which prior interface geometries are desired to be recapitulated in the samples.

The resulting model, `cc83`, has the same architecture as `cc58`, except for one additional input channel for the boolean hotspot mask, and was trained for 1.25M steps. Despite not being trained on protein complexes with more than two chains, the model is capable of conditioning complex generation on two or three target chains. Figure 8 shows the SpCas9 target in BindCraft which was modeled as three chains due to the two chain breaks in the target structure. The structure shown has iPTM = 0.76, iPAE = 0.26, complex pLDDT = 0.86, and binder scRMSD = 0.95 Å using AF2 single-sequence model 1.

**Figure 8:**
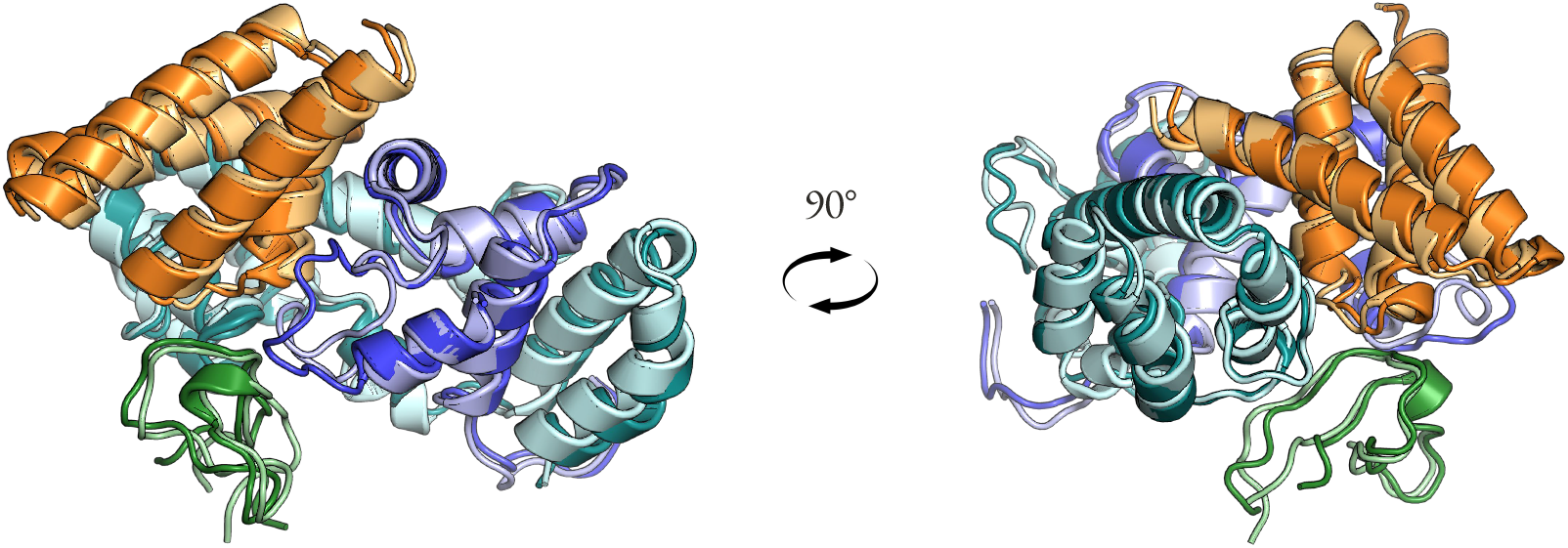
Generalization to more than two chains. SpCas9 target in BindCraft is modeled as three chains (blue/cyan/green) and the generated binder is the fourth chain (orange). Darker shades are chains from the structure predicted by AF2 single-sequence and lighter shades are chains from the model `cc83` after PyRosetta relax.

We evaluated `cc83` on the BindCraft targets using the following set of AlphaFold2-based thresholds, following BindCraft: complex pLDDT *>* 0.80, iPTM *>* 0.5, iPAE < 0.35, and *apo* binder RMSD < 3.5 Å. We note that a direct comparison may be misleading as `cc83` is a base model which was trained on protein-protein interfaces not subject to filtering by AF2 metrics or sampling-time guidance. In contrast, BindCraft explicitly optimizes for AF2 pLDDT, iPAE, and iPTM, though it uses the multimer model for design and the monomer model for evaluation. To be as close as possible to BindCraft’s evaluation pipeline, we designed the sequence using AF2-Multimer hallucination with binder template unmasked but interchain template features masked. Both sequence and sidechain features of the target were given during design. All other design parameters were identical to BindCraft’s four-stage hallucination protocol. PyRosetta then determined the interface residues and these remain fixed positions during ProteinMPNN sequence redesign. For evaluation, we follow BindCraft and use AF2 in single-sequence mode, providing the template of the target, no template of the binder, remove interchain template features, and do not use initial guess. However, we remove the target template sequence features to allow for target backbone flexibility. This is due to the finding that many BindCraft designs would not pass the iPTM and iPAE thresholds on a subtly different target conformation if the target sequence was left unmasked.

To compare with BindCraft runs, we took a random subsample of 100 passing BindCraft trajectories (non-clashing, not low confidence) per target and took the better sequence from the first two ProteinMPNN redesigned sequences by AF2 single-sequence metrics. The results are shown in Figure 9. By AF2 metric-based success rates, BindCraft outperforms Protpardelle-1c on all targets except DerF21. We note that the comparison is not one-to-one, as we drew generally much fewer samples and do not apply any filtering to the Protpardelle-1c samples, nor compare against low confidence and clashing trajectories which are discarded by BindCraft during hallucination. Another distinction is the abundance of alpha-helical samples generated by BindCraft relative to Protpardelle-1c (Figure 10). BindCraft directly optimizes for sequences favored by AF2, while Protpardelle-1c was a base model not optimized towards any specific metric. As shown in SHAPES [16], helical structures contribute to the bias towards *in silico* designability: beta-containing structures are more difficult for AF2 single-sequence, whereas helices are more favored and easier to predict. It is unclear whether the Protpardelle-1c samples are indeed difficult to pass *in silico* filters due to AF2 preference for helical structures. Nonetheless, the nonzero samples from 100 random models that pass the AF2 metrics suggest potential future improvements using reward-based guidance and alternative binder sequence design methods. Experimental testing may also be necessary to validate some plausible but non-helical samples that fail AF2 filters.

**Figure 9:**
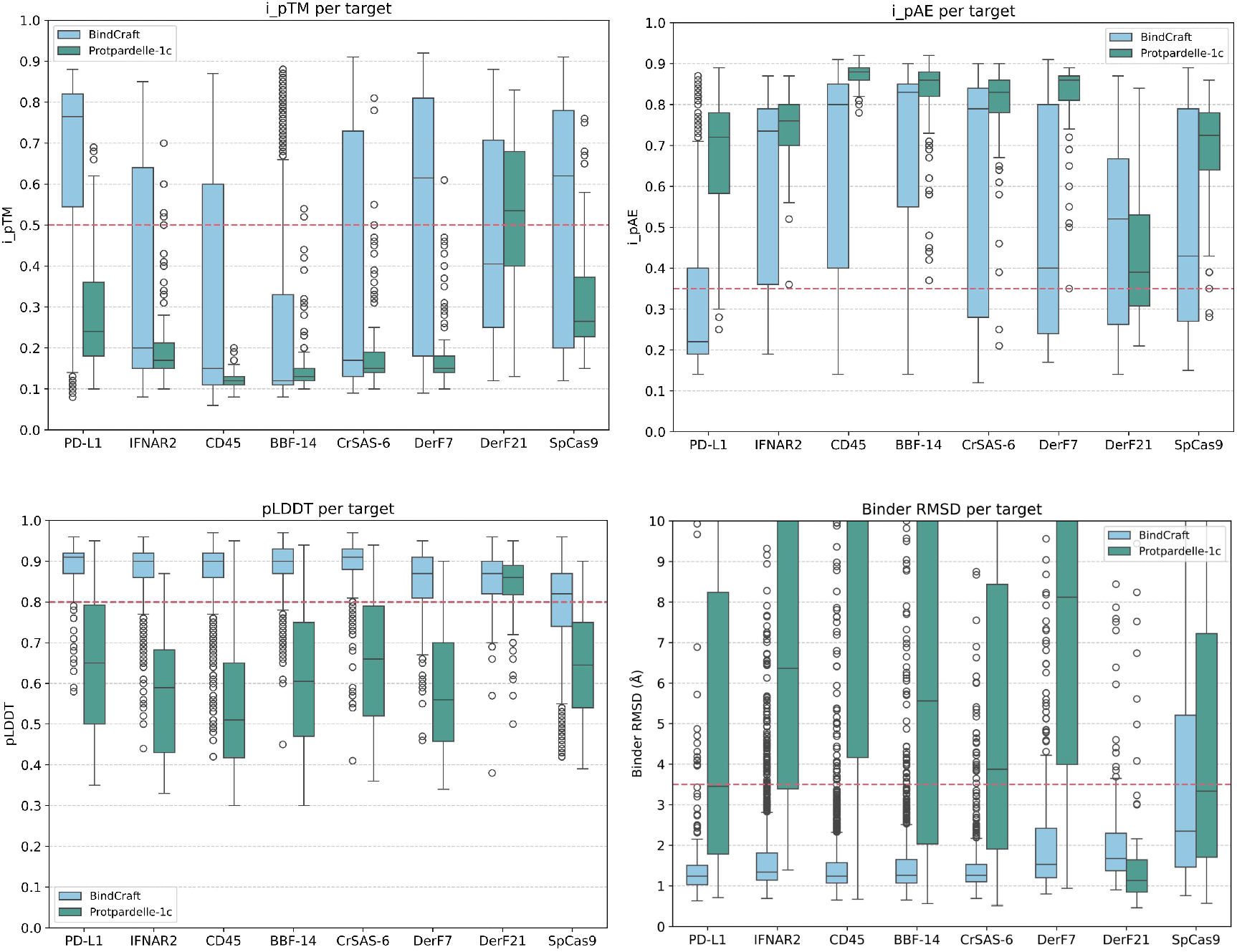
Protpardelle-1c cc83 vs. BindCraft. 100 samples from Protpardelle-1c cc83 compared against ProteinMPNN-reoptimized sequences from 100 subsampled BindCraft trajectories. The success thresholds are denoted as dashed lines: iPTM > 0.5, iPAE < 0.35, pLDDT > 0.8, binder RMSD < 3.5 Å.

**Figure 10:**
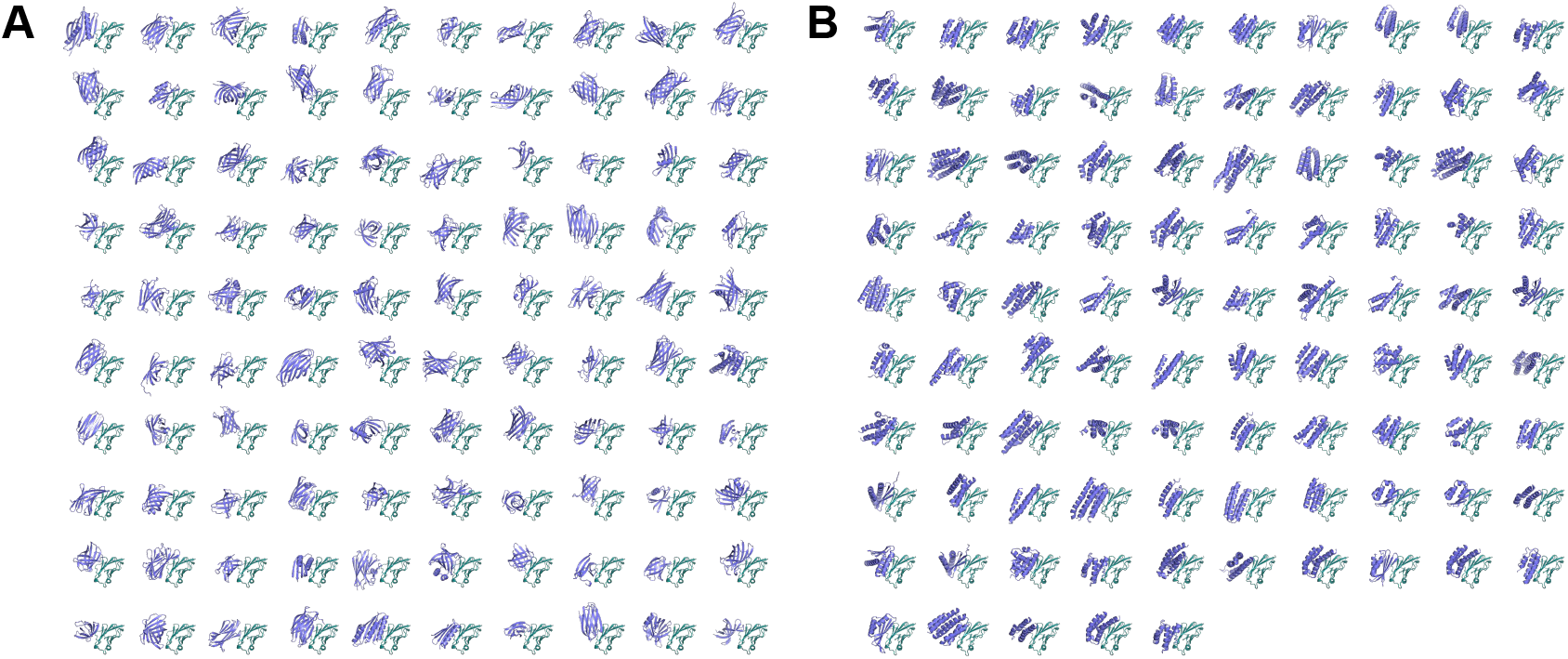
BindCraft binders are more enriched in helices. (A) 100 random samples from `cc83` given PD-L1 as the target. (B) AF2 single-sequence model 1 structures of 95 BindCraft samples on PD-L1 which pass all BindCraft *in silico* filters.

## 4 Discussion

Protpardelle-1c is an update to the original Protpardelle where the main differences are the self-consistent CATH training data and a more light-weight noise level embedding mechanism. The most impactful change to training stability was the appropriate handling of unresolved residues. The recommended model is `cc58` for backbone-only single-chain conditional generation, `cc83` for backbone-only multi-chain conditional generation, `cc89` for all-atom single-chain structure refinement, and `cc91` for all-atom single-chain conditional generation. Additional models are detailed in Table 2.

We only evaluated the final model checkpoint and have not evaluated auto-guidance [21], where an earlier checkpoint is used as a bad version to guide against, post-hoc EMA [22], or sweeped classifier-free guidance scales. Feynman-Kac steering using sequence design model likelihood may also be promising [23]. The training data was aggressively filtered to retain examples with scRMSD < 2.0 Å and pLDDT *>* 80; we anticipate that scaffold diversity can be further improved by training on medium to low confidence structures at medium to high noise levels, respectively, as in Ambient Protein Diffusion [24]. We also anticipate that a similar filter applied to the chain pair dataset may lead to improved performance on the BindCraft benchmark.

For motif scaffolding, several comparisons remain: a baseline for searching motifs in a large database of native structures (e.g. AlphaFold Database [25]) or sampled structures (e.g. SHAPES [16]) and comparison to recent diffusion models beyond RFdiffusion. The parameters which control how motifs are generated on the fly can also be tuned, in particular, by aligning the types of motifs generated during training with features commonly observed in functional motifs, e.g. substructures more enriched in loops, may further improve performance. Analysis of the frequencies of motif structures observed during training compared to the scaffolding success of structurally similar motifs may be insightful in explaining the heterogeneity of motif scaffolding problem difficulty. A limitation of current motif scaffolding benchmarks is that the occlusion of function sites is not considered in the criteria. For experimental applications, filtering out samples where the function motif is occluded would reduce success rates and additional guidance terms to prevent clashes may be necessary.

## 5 Acknowledgments

We thank Braxton Bell, Hyejin Lee, and Paul Ruijgrok for feedback on the codebase. We thank Brian Trippe, Wei Deng, and Christian Choe for helpful discussions. We thank Martin Pacesa for providing raw BindCraft trajectories. We thank Alex Chu for the original Protpardelle codebase. The computing for this project was performed on the Sherlock cluster. We would like to thank Stanford University and the Stanford Research Computing Center for providing computational resources and support that contributed to these research results. Petr Kouba acknowledges the computational resources provided at the LUMI supercomputer owned by the EuroHPC Joint Undertaking, hosted by CSC (Finland) with access granted through IT4I National Supercomputing Center, Czech Republic and the e-INFRA CZ (No. 90254) project. The research stay of Petr Kouba at Stanford was funded by the European projects CLARA (No. 101136607), ERC project FRONTIER (No. 101097822), COST Action CA21162 COZYME as well as by G-Research, Loschmidt Laboratories, Czech Technical University in Prague and Masaryk University. Petr Kouba thanks Jiří Damborský, Josef Šivic and Stanislav Mazurenko for their support. T.L. and Z.L. are supported by Stanford Graduate Fellowship. R.W.S acknowledges funding support from the NSF Graduate Research Fellowship (DGE-2146755). This project is supported by NIH (R01GM147893 to P.-S.H.), Merck Research Laboratories (MRL) Scientific Engagement and Emerging Discovery Science (SEEDS) Program, and Stanford Medicine Catalyst. The views and conclusions contained in this document are those of the authors and should not be interpreted as representing the official policies, either expressed or implied, of the U.S. Government.

## Appendix A Chain pair dataset

For training multi-chain models, we extracted a dataset of chain pairs involved in a protein-protein interface from the PDB using Boltz-1 data pre-processing scripts [20]. Following AlphaFold3, we defined a pair of chains to be involved in an interface if any heavy atoms on one chain were within 5Å of another chain [26]. To define interface clusters, we first clustered all protein chains using MMseqs2 [27] at a 40% sequence identity threshold. Similar to AlphaFold3, we defined two interfaces as part of the same cluster if both chains in each interface corresponded to the same pair of chain clusters. We then filtered for structures with a resolution of 2Å or less and for interfaces where each chain contained at least 32 residues and at most 256 residues, yielding 1,593,738 interfaces. During training, a random cluster is sampled then a random member is sampled from each cluster. We use each `.cif` file’s `_atom_site.label_seq_id` to derive positional encodings.

## Appendix B mdCATH-based dataset

### Cluster-based subsampling of Molecular Dynamics trajectories

We construct the dataset for the training of Protpardelle by sampling 32 conformations per protein out of the respective conformational ensembles from mdCATH. To sample the structures, we perform k-median clustering (*k* ∈ {2, 3, 4, 5}) of the conformational ensembles in a 5-dimensional projection of the coordinate space. The projection was obtained using Variational Approach to Markov Processes (VAMP) [28], a dimensionality reduction technique which unlike PCA and similarly to TICA takes into account not only the covariances of the coordinates at a given timepoint, but it also considers the time-lagged covariances between structures from different timepoints separated by a fixed lagtime *τ* (in our case *τ* ∈ {1*ns*, 5*ns*, 10*ns*, 15*ns*, 20*ns*}) assuming the Markovianity of the underlying process. To settle on particular clusterings, we selected the optimal hyperparameters *k* and *τ* using Silhouette rule [30] for each protein. From each of the *k* clusters we then drew 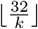 samples, including the cluster center and 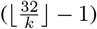 uniform samples from each cluster. The remaining (32 mod *k*) samples were sampled uniformly from randomly selected clusters.

### Fixing bond geometry of MD samples

The MD simulation protocol of mdCATH did not enforce the correct bond lengths and bond angles, therefore such geometry might be distorted in the conformations sampled directly from the MD trajectories. To correct for this, we performed Rosetta cartesian minimization. To keep as much conformational diversity as possible, we constrained the C*α* coordinates with a harmonic potential.

